# Hypothalamic MCH neurons links tau pathology to sleep disruption

**DOI:** 10.64898/2026.05.31.729026

**Authors:** Ke Pang, Siyin Guo, Ruilin Yang, Ajay Kumar, Wyatt D. Morse, Leann Xu, Qiaoling Liu, Dequan Jiang, Peng Zhong

## Abstract

Sleep disruption is an early and pervasive feature of Alzheimer’s disease (AD), yet the circuit mechanisms linking tau pathology to sleep-wake dysfunction remain unresolved. Here, we identify melanin-concentrating hormone (MCH) neurons in the lateral hypothalamus (LH) as a critical node disrupted in tauopathy. Longitudinal EEG/EMG recordings in *PS19* mice reveal progressive impairments in sleep architecture and homeostasis. Histological analyses revealed significant degeneration of both MCH neurons and the neighboring hypocretin (Hcrt) neuronal population in the LH at late stages of tauopathy. *In vivo* fiber photometry recordings demonstrated a selective functional impairment of MCH neurons, characterized by reduced activity during REM sleep, whereas Hcrt neuronal activity remained largely preserved. In addition, cell-autonomous expression of mutant tau in MCH neurons recapitulates sleep disruption, establishing a causal role. Despite tauopathy-induced neuronal loss and reduced endogenous activity, optogenetic and chemogenetic activation show that MCH neurons retain functional capacity, and their activation restores sleep in aged *PS19* mice. Together, these findings define a circuit mechanism linking tau pathology to sleep and identify MCH neurons as a tractable therapeutic target.

## INTRODUCTION

Sleep disturbances are among the earliest and most prevalent non-cognitive symptoms of Alzheimer’s disease (AD) (*1, 2*), affecting up to two-thirds of patients and substantially contributing to disease burden and institutionalization (*2–4*). Alterations in sleep architecture, including reduced sleep efficiency, increased nighttime awakenings, excessive daytime sleepiness, and rapid eye movement (REM) sleep abnormalities, often precede cognitive decline (*5–7*) and worsen as the disease progresses (*7*). Growing evidence suggests that sleep disruption is not merely a consequence of neurodegeneration but actively contributes to AD pathogenesis. A defining pathological hallmark of AD, frontotemporal dementia (FTD), and other neurodegenerative tauopathies is the hyperphosphorylation and aggregation of the microtubule-associated protein tau into neurofibrillary tangles (NFTs) (*8, 9*), which are closely associated with synaptic dysfunction, neuronal loss, progressive cognitive decline, and disease severity (*3, 10, 11*). Experimental sleep deprivation increases extracellular tau levels and accelerates tau pathology, whereas sleep promotes the clearance of neurotoxic proteins from the brain (*12*). These observations support a bidirectional relationship between tau pathology and sleep disruptions seen in related neurodegenerative disorders.

Tau pathology accumulates early in multiple brain regions involved in sleep-wake regulation, including the locus coeruleus, dorsal raphe, basal forebrain, tuberomammillary nucleus, and lateral hypothalamus (LH) (*3*). Consistent with these neuropathological findings, transgenic mouse models expressing human tau develop sleep disturbances resembling those observed in AD patients and other human tauopathies (*12–15*). However, the neural circuit mechanisms through which tau pathology disrupts sleep remain poorly understood. Identifying the specific neuronal populations affected by tau pathology is essential for understanding the origins of sleep dysfunction and developing targeted therapeutic strategies.

The LH contains two anatomically intermingled but functionally distinct neuronal populations that play critical roles in sleep-wake regulation (*16*): hypocretin/orexin (Hcrt) neurons and melanin-concentrating hormone (MCH) neurons. Hcrt neurons are essential for maintaining wakefulness and behavioral arousal (*17–21*), whereas MCH neurons are preferentially active during REM sleep (*22*), respond to REM sleep deprivation and integrate homeostatic sleep drive (*23–26*), and mainly promote and maintain REM-phase of sleep (*22, 27–30*). MCH neurons are particularly intriguing in the context of AD because REM sleep abnormalities are strongly associated with cognitive decline (*31, 32*), and MCH signaling influences both sleep and hippocampal-dependent memory (*33*). Moreover, NFTs (*34, 35*) and neuronal loss (*34, 36*) have been reported in the LH of AD patients, and cerebrospinal fluid MCH levels correlate with tau burden, REM sleep disruption, and cognitive impairment (*37*). Despite these observations, it remains unknown whether tau pathology directly impairs MCH and Hcrt neuronal function and whether such dysfunction contributes to sleep disturbances in tauopathy.

Here, we combined longitudinal EEG/EMG recordings, histological analyses, *in vivo* fiber photometry, and causal circuit manipulations to determine how tau pathology affects LH sleep/wake-regulatory neurons in mouse models of tauopathy. We show that tauopathy produces progressive sleep disturbances accompanied by degeneration of both MCH and Hcrt neurons, but selectively impairs REM sleep-associated activity of MCH neurons. Remarkably, despite neuronal loss and functional deficits, the remaining MCH neurons retain their sleep-promoting capacity. Optogenetic and chemogenetic activation of MCH neurons restores sleep quantity and quality in tauopathy mice, identifying MCH circuit dysfunction as a contributor to disease-associated sleep disturbances and highlighting the MCH system as a potential therapeutic target for treating sleep dysfunction in AD and related tauopathies.

## RESULTS

### Sleep and wake are disrupted in mouse stains of tauopathy

To determine the temporal emergence of sleep abnormalities during tauopathy progression, we longitudinally assessed sleep architecture in *PS19* mice and wild-type (WT) littermates using chronic ectroencephalogram (EEG)/electromyogram (EMG) recordings. 24-hour baseline sleep recordings were collected collected repeatedly from the same cohort across multiple ages to track age-dependent changes in sleep-wake regulation (Fig. 1). Wakefulness, REM, and NREM states were classified based on EEG/EMG recordings. There were no detectable differences in total wakefulness, NREM sleep, REM sleep, or bout architecture at 8 months of age between genotypes (Fig. 1A). In contrast, by 10-11 months of age, *PS19 mice* exhibited marked alterations in sleep organization. Relative to WT littermates, *PS19 mice* spent significantly more time awake and less time in both NREM and REM sleep across the recording period (Figs. 1 E and G). These changes were accompanied by fragmentation of sleep states, characterized by prolonged wake bouts and shortened NREM and REM sleep bouts (Figs. 1F and H), indicating reduced stability of sleep maintenance rather than simple redistribution of vigilance states.

**Figure 1.**
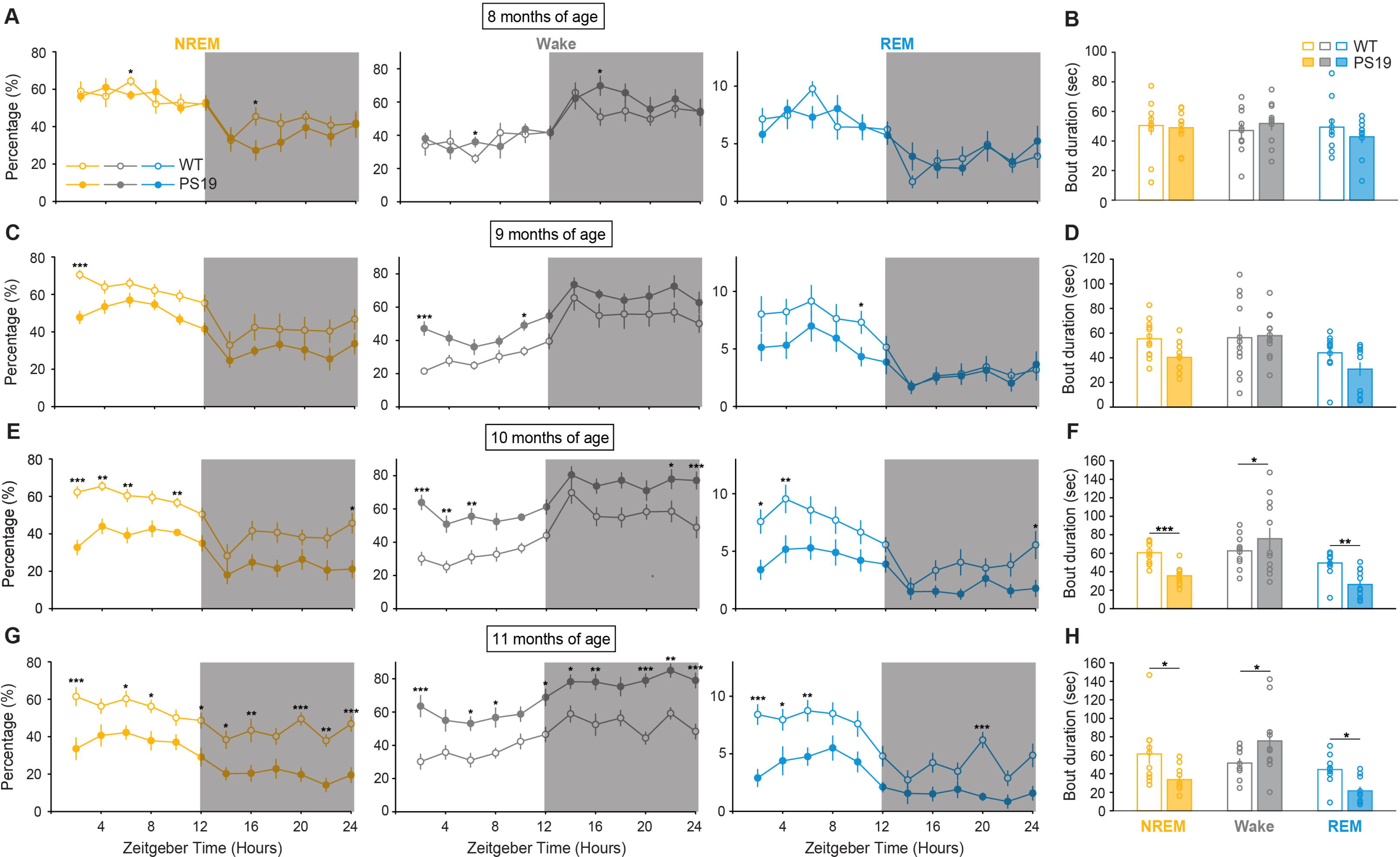
Age-dependent alterations in sleep-wake architecture and bout duration in *PS19* mice. (A) Percentage of time spent in NREM sleep, wake, and REM sleep across the 24-h light/dark cycle in 8-month-old WT and *PS19* mice. Data are plotted as a function of Zeitgeber time. Gray shading indicates the dark phase. Yellow, NREM; gray, wake; blue, REM. Open circles, WT; filled circles, *PS19*. n = 9-12 mice per group; Error bars, ±SEM. (B) Quantification of mean bout duration for NREM sleep, wake, and REM sleep in 8-month-old WT and *PS19* mice. Each circle represents one mouse. Bars, mean ± SEM. (C) Similar to (A), for 9-month-old WT and *PS19* mice. (D) Similar to (B), for 9-month-old WT and *PS19* mice. (E) Similar to (A), for 10-month-old WT and *PS19* mice. *PS19* mice showed reduced NREM sleep during the light phase and increased wakefulness compared with WT mice. (F) Similar to (B), for 10-month-old WT and *PS19* mice. *PS19* mice showed shorter NREM and REM bout durations and longer wake bout duration compared with WT mice. (G) Similar to (A), for 11-month-old WT and *PS19* mice. *PS19* mice showed persistent reduction in NREM sleep and increased wakefulness across multiple Zeitgeber time points, together with reduced REM sleep at selected time points. (H) Similar to (B), for 11-month-old WT and *PS19* mice. *PS19* mice showed shortened NREM and REM bout durations and prolonged wake bout duration compared with WT mice. Data are presented as mean ± SEM. For time-course analyses of sleep-wake state percentage, statistical significance was determined by two-way ANOVA followed by post hoc multiple-comparisons tests. For comparisons of bout duration between treatment groups within each vigilance state, Welch’s t-test was used. Asterisks indicate significant differences between WT and PS19 mice at individual Zeitgeber time points or within each behavioral state. *P < 0.05; **P < 0.01; ***P < 0.001.

Spectral analysis of EEG activity was performed across all epochs classified as wake, NREM and REM sleep (Fig. 2). Figure 2A shows representative 24-hour EEG spectrograms from individual WT and *PS19* mouse at 10 months of age. Compared with age-matched WT littermate, *PS19* mouse exhibited reduced EEG spectral power in the low-frequency 0-5 Hz range, particularly during the dark phase. Two-way ANOVA revealed significant genotype × frequency band interactions at 10 months of age across all vigilance states (two-way ANOVA, P < 0.0001 for all brain states; Fig. 2F). EEG power spectra were quantified across predefined frequency bands, including delta (δ, 0.5-4 Hz), low theta (Lθ, 4-8 Hz), high theta (Hθ, 8-10 Hz), low gamma (Lγ, 15-50 Hz), high gamma (Hγ, 60-80 Hz), and very high gamma (Vhγ, 90-200 Hz). Compared with WT littermates, *PS19* mice exhibited selective alterations in θ-band activity during disease progression. Specifically, power in the Lθ band during wakefulness was significantly elevated in *PS19* mice at 8-10 months of age (P < 0.001, two-tailed *t*-test; Figs. 2C, E and G), whereas Lθ power during REM sleep was significantly reduced at 11 months of age (P < 0.01; Fig. 2F). These findings indicate state-dependent disruption of cortical oscillatory dynamics in *PS19* mice, with enhanced low-frequency activity during wakefulness and diminished REM-associated theta oscillations emerging at later stages of tauopathy.

**Figure 2.**
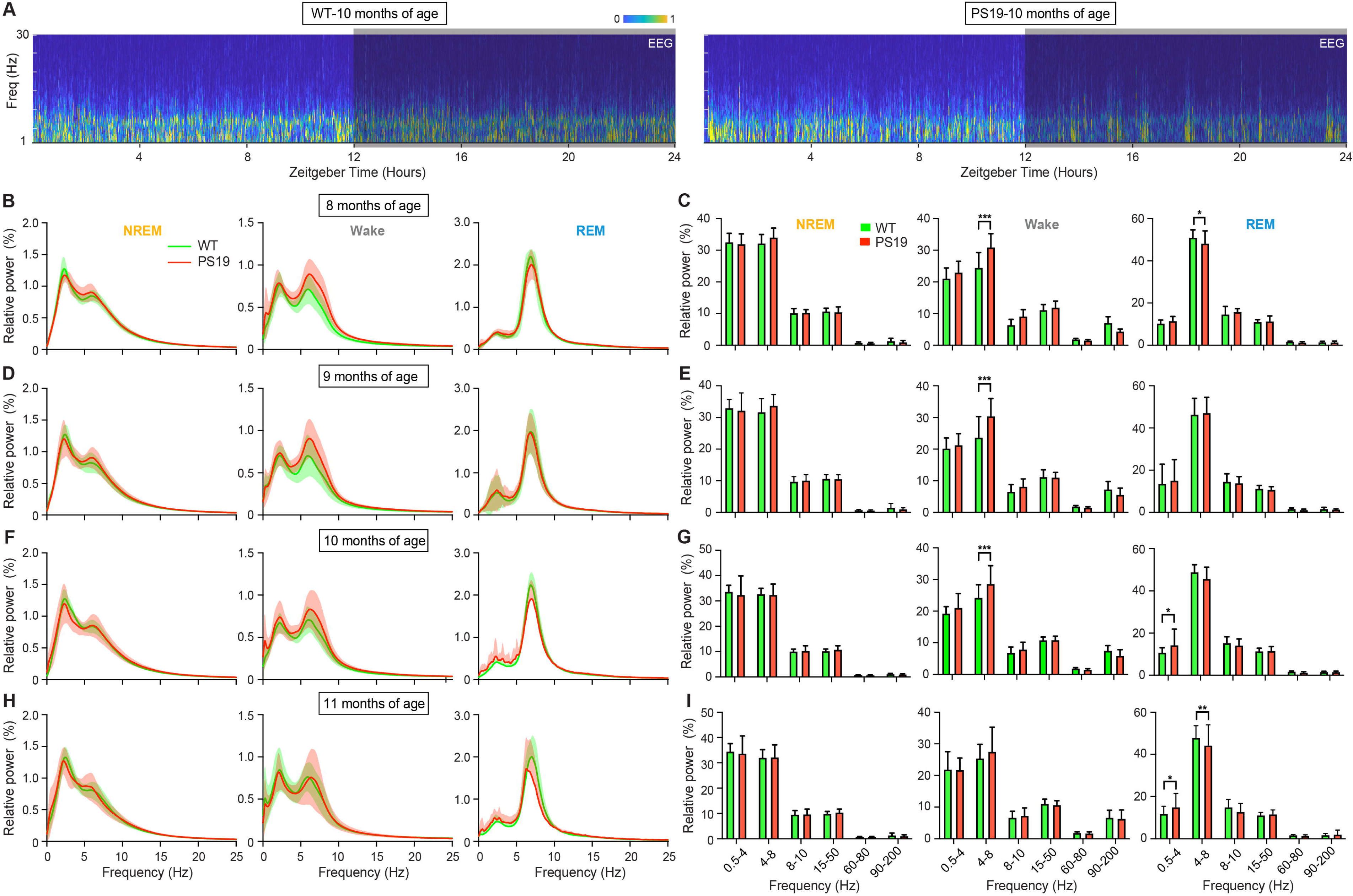
Age-dependent EEG spectral alterations across vigilance states in *PS19* mice. (A) Example EEG power spectrograms recorded over 24 h from WT and *PS19* mice at 10 months of age. The x-axis indicates Zeitgeber time, and the y-axis indicates EEG frequency. Gray shading indicates the dark phase. Color scale represents normalized EEG power. (B) Relative EEG power spectra during NREM sleep, wake, and REM sleep in 8-month-old WT and *PS19* mice. Green, WT; red, PS19. Shading, ±SEM. (C) Quantification of EEG relative power in the indicated frequency bands during NREM sleep, wake, and REM sleep in 8-month-old WT and *PS19* mice. Bars, mean ± SEM. (D and E) Similar to (B) and (C), for 9-month-old WT and *PS19* mice. (F and G) Similar to (B) and (C), for 10-month-old WT and *PS19* mice. (H and I) Similar to (B) and (C), for 11-month-old WT and *PS19* mice. Asterisks indicate significant differences between WT and *PS19* mice. Statistical significance for relative EEG power spectra and frequency-band quantification was determined by two-way ANOVA followed by post hoc multiple-comparisons tests. *P < 0.05; **P < 0.01; ***P < 0.001.

To determine whether sleep abnormalities extend beyond the *PS19* model and represent a broader feature of tauopathy, we additionally examined sleep-wake regulation in *hTau* mice (Fig. 3), an independent mouse model that develops progressive tau pathology. *hTau* mice exhibit age-dependent pathological tau accumulation characterized by somatodendritic redistribution of tau, tau hyperphosphorylation, accumulation of paired helical filaments, and the formation of NFTs(*38*). Unlike *PS19* mice, *hTau* mice showed no significant differences from WT littermates in total time spent in all brain states between 9 and 12 months of age (Figs. 3A, C, E, and G). Sleep architecture remained largely preserved, as NREM and REM sleep bout durations were comparable between genotypes (Figs. 3B, D, F, and H). Interestingly, wake bout duration was significantly reduced in *hTau* mice at 10 and 11 months of age relative to WT controls (Figs. 3D and F), suggesting altered wake-state maintenance preceding overt changes in overall sleep time. Spectral analysis revealed progressive and state-dependent changes in oscillatory dynamics in *hTau* mice (Fig. 4). During NREM sleep, *hTau* mice exhibited reduced δ power, with the effect becoming progressively greater from 10 to 12 months of age (Figs. 4C-H). Together, these findings indicate that sleep dysfunction in *hTau* mice emerges later and manifests differently than in *PS19* mice, with selective vulnerability of REM sleep and wake consolidation accompanied by impaired NREM-associated cortical oscillatory activity.

**Figure 3.**
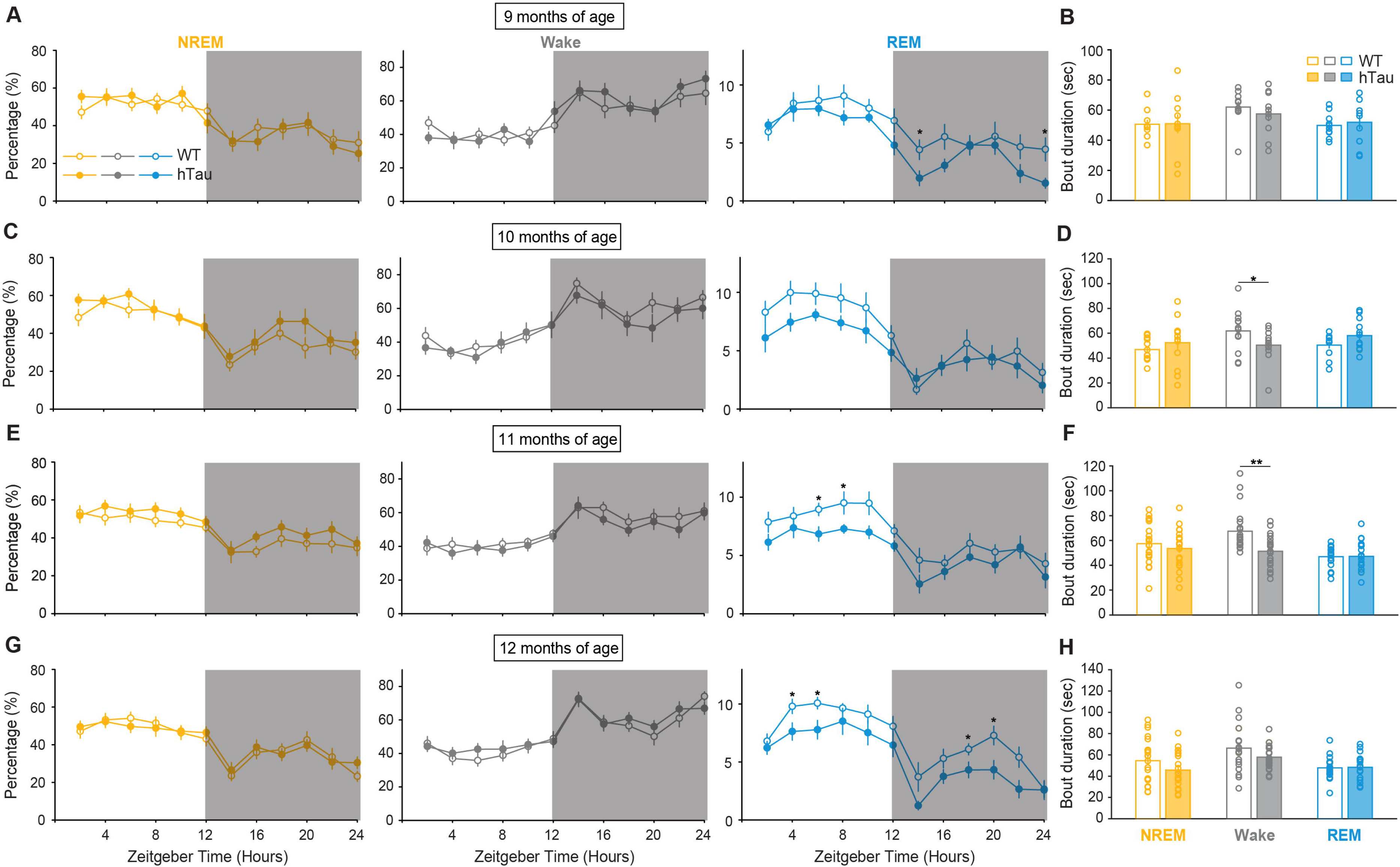
Sleep-wake architecture and bout duration in aging *hTau* mice. (A) Percentage of time spent in NREM sleep, wake, and REM sleep across the 24-h light/dark cycle in 9-month-old WT and hTau mice. Gray shading indicates the dark phase. Yellow, NREM; gray, wake; blue, REM. Open circles, WT; filled circles, *hTau*. N = 11-13 mice per group; Error bars, ±SEM. (B) Quantification of mean bout duration for NREM sleep, wake, and REM sleep in 9-month-old WT and *hTau* mice. Each circle represents one mouse. Bars, mean ± SEM. (C and D) Similar to (A) and (B), for 10-month-old WT and *hTau* mice. (E and F) Similar to (A) and (B), for 11-month-old WT and *hTau* mice. (G and H) Similar to (A) and (B), for 12-month-old WT and hTau mice. *hTau* mice displayed no major differences in the percentage of time spent in NREM sleep, wake, or REM sleep across the 24-h recording period, but showed reduced wake bout duration at 10 and 11 months of age. Statistical significance for sleep-wake state percentage in (A, C, E, and G) was determined by two-way ANOVA followed by post hoc multiple-comparisons tests. Statistical significance for mean bout duration in (B, D, F, and H) was determined by Welch’s t-test between WT and hTau mice within each vigilance state. *P < 0.05; **P < 0.01.

**Figure 4.**
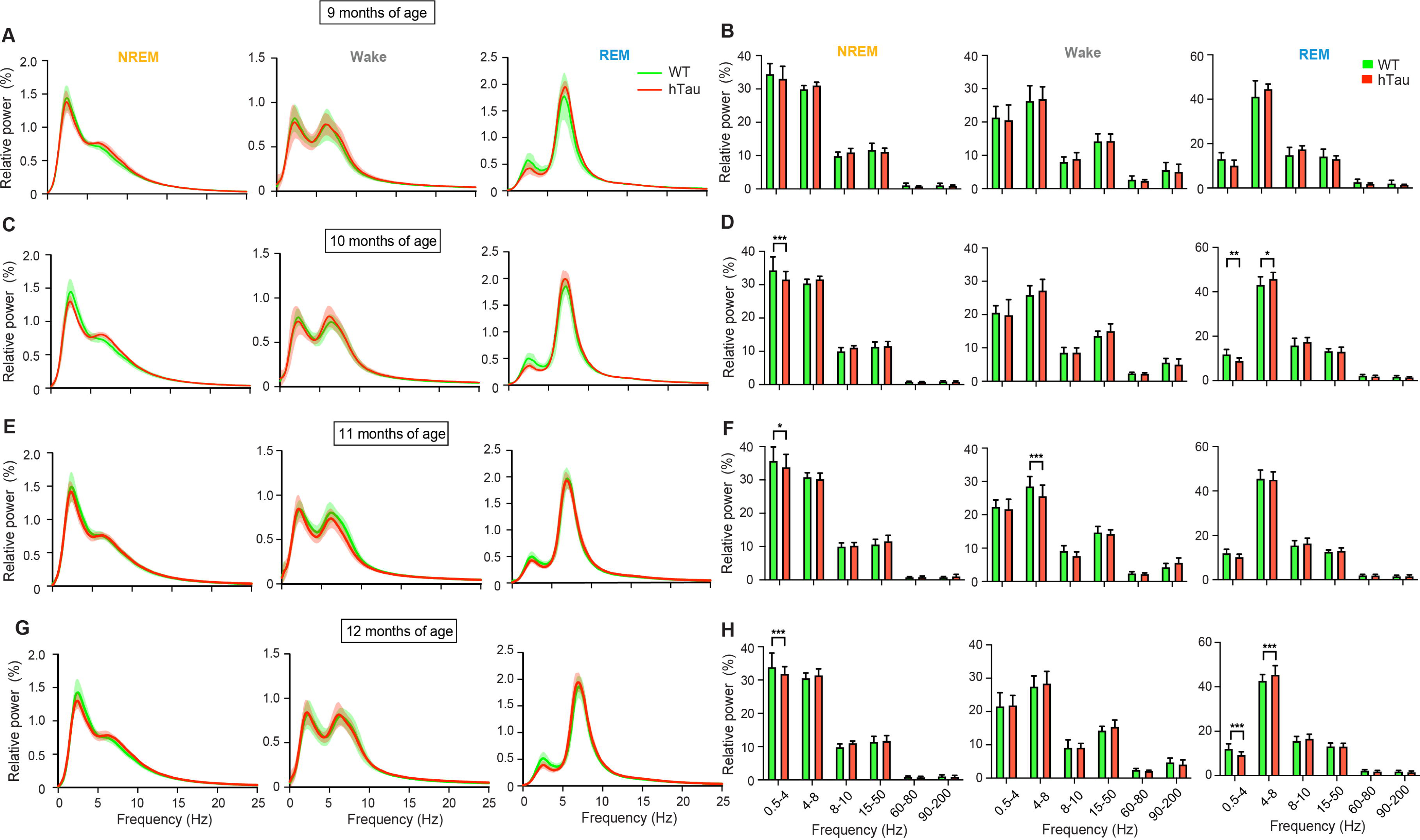
Age-dependent EEG spectral alterations across vigilance states in *hTau* mice. (A) Relative EEG power spectra during NREM sleep, wake, and REM sleep in 9-month-old WT and *hTau* mice. Green, WT; red, *hTau*. Shading, ±SEM. (B) Quantification of EEG relative power in the indicated frequency bands during NREM sleep, wake, and REM sleep in 9-month-old WT and *hTau* mice. Bars, mean ±SEM. (C and D) Similar to (A) and (B), for 10-month-old WT and *hTau* mice. (E and F) Similar to (A) and (B), for 11-month-old WT and *hTau* mice. (G and H) Similar to (A) and (B), for 12-month-old WT and *hTau* mice. *hTau* mice showed age-dependent, frequency-band-specific changes in EEG spectral power across vigilance states. Asterisks indicate significant differences between WT and hTau mice. Statistical significance for relative EEG power spectra in (A, C, E, and G) and frequency-band quantification in (B, D, F, and H) was determined by two-way ANOVA followed by post hoc multiple-comparisons tests. *P < 0.05; **P < 0.01; ***P < 0.001.

Because sleep homeostasis is a fundamental mechanism that restores sleep following prolonged wakefulness, we next evaluated the ability of *PS19* and *hTau* mice to mount compensatory rebound sleep after acute sleep loss. Beginning at light onset (ZT0), 11-month-old *PS19* mice, 12-month-old *hTau* mice, and their respective age-matched WT littermates underwent 5 h of sleep deprivation (SD), during which they were kept awake by the introduction of novel objects or gentle tapping on the cages(*39*). Mice were subsequently allowed undisturbed recovery sleep (RS). Compared with WT littermates, *PS19* mice exhibited marked reductions in NREM sleep during the recovery period following 5 h SD (Fig. 5A), whereas *hTau* mice showed only modest impairments in REM sleep recovery (Fig. 5B).

**Figure 5.**
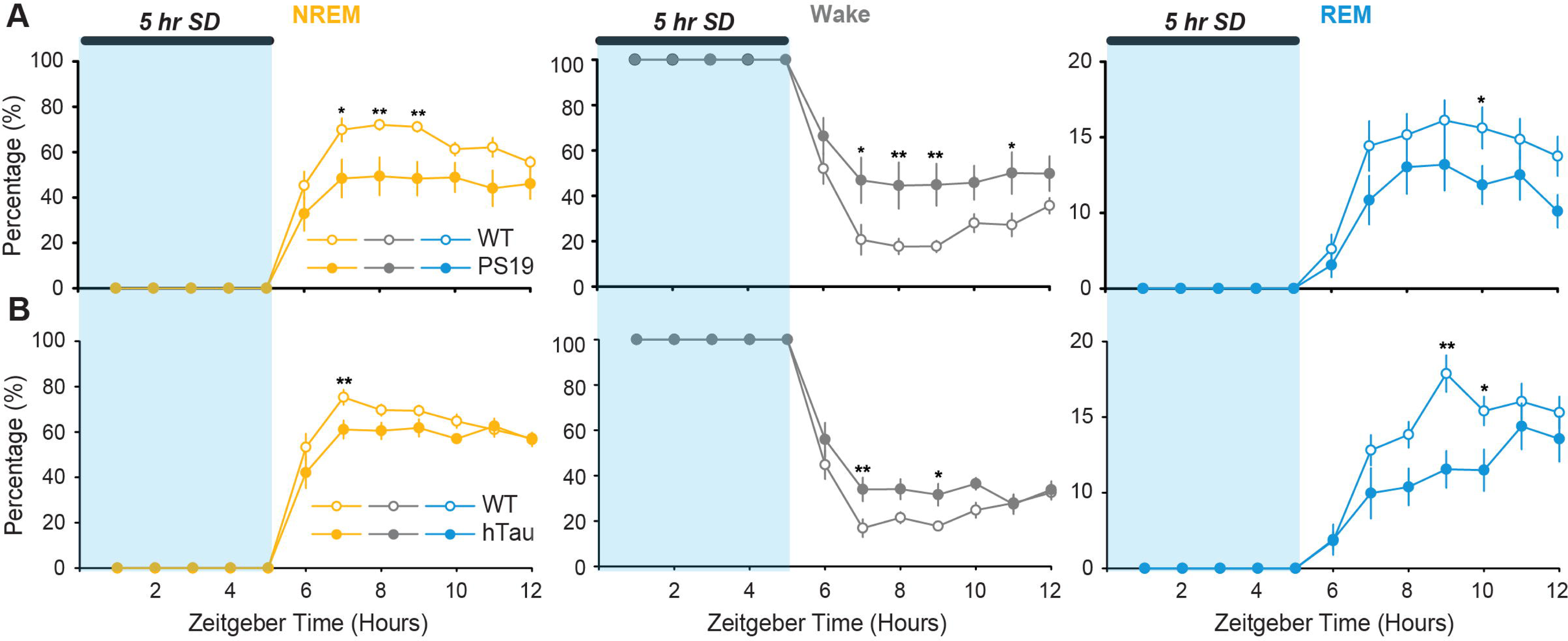
*PS19* and *hTau* mice exhibit impaired sleep homeostatic recovery following 5-h sleep deprivation. (A) Percentage of time spent in NREM sleep, wake, and REM sleep during and after 5-h sleep deprivation in WT and *PS19* mice at 11 months of age. Blue shading indicates the 5-h sleep deprivation period. Data are plotted as a function of Zeitgeber time. Yellow, NREM; gray, wake; blue, REM. Open circles, WT; filled circles, *PS19*. n = 8 mice per group; Error bars, ±SEM. (B) Percentage of time spent in NREM sleep, wake, and REM sleep during and after 5-h sleep deprivation in WT and *hTau* mice at 12 months of age. Blue shading indicates the 5-h sleep deprivation period. Yellow, NREM; gray, wake; blue, REM. Open circles, WT; filled circles, *hTau*. n = 11-12 mice per group; Error bars, ±SEM. Following sleep deprivation, *PS19* and *hTau* mice showed altered sleep rebound responses compared with WT mice, including reduced NREM recovery and increased wakefulness during the rebound period, with genotype-dependent changes in REM sleep recovery. Asterisks indicate significant differences between WT and transgenic mice at individual Zeitgeber time points. Statistical significance was determined by two-way ANOVA followed by post hoc multiple-comparisons tests. *P < 0.05; **P < 0.01.

### Selective loss of MCH and Hcrt neurons emerges at late stages of tauopathy

Pathological tau accumulation was abundant in the lateral hypothalamus (LH) of *PS19* mice at late stages of tauopathy, a region that contains two key sleep–wake regulatory neuronal populations: melanin-concentrating hormone (MCH) neurons and hypocretin (Hcrt) neurons. These intermingled LH neuronal populations project broadly throughout the brain and play critical roles in sleep–wake regulation(*18, 40*). We therefore quantified the neuronal density of MCH and Hcrt neurons in 11-month-old PS19 mice exhibiting sleep disturbances and found significant reductions in both neuronal populations compared with WT controls (Fig. 6), indicating that these LH sleep/wake neurons are highly vulnerable under tauopathy conditions.

**Figure 6.**
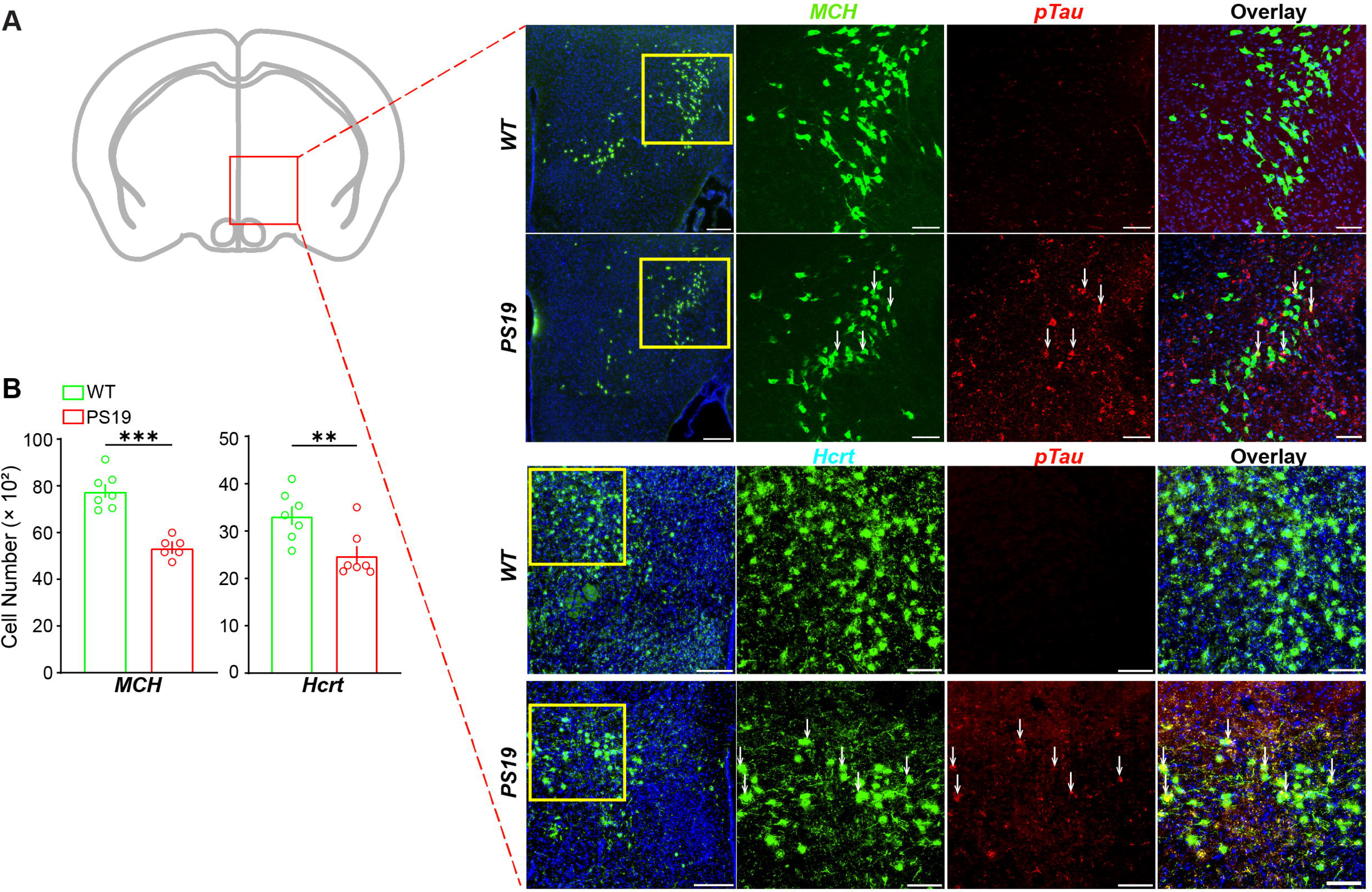
*PS19* mice exhibit phosphorylated tau accumulation and loss of MCH and Hcrt neurons in the lateral hypothalamus. (A) Left, schematic showing the coronal brain section and the lateral hypothalamic region analyzed in this study. Red box indicates the region used for immunofluorescence imaging. Right, representative fluorescence images from WT and *PS19* mice showing MCH-positive neurons or Hcrt-positive neurons together with phosphorylated tau staining in the lateral hypothalamus. For the upper panels, green indicates MCH, red indicates pTau (AT-8), and blue indicates nuclear counterstain. For the lower panels, green indicates Hcrt, red indicates pTau, and blue indicates nuclear counterstain. Yellow boxes indicate regions shown at higher magnification in the adjacent single-channel and merged images. Arrows indicate pTau-positive signals in *PS19* mice within regions containing MCH- or Hcrt-positive neurons. Overlay images show the spatial relationship between pTau signals and MCH- or Hcrt-positive neuronal populations. Scale bars, 200 μm for overview images and 100 μm for enlarged images. (B) Quantification of MCH-positive and Hcrt-positive cell numbers in the lateral hypothalamus of WT and *PS19* mice. Each circle represents one sample. n = 6-7 mice per group; Bars, mean ± SEM. *PS19* mice showed a significant reduction in both MCH-positive and Hcrt-positive cell numbers compared with WT mice. Statistical significance was determined by Welch’s t-test. **P < 0.01; ***P < 0.001.

### Aberrant activity of lateral hypothalamic sleep/wake neurons at late stages of tauopathy

To investigate whether tauopathy alters the activity of hypothalamic sleep–wake neuronal populations, we performed fiber photometry recordings of GCaMP signals from lateral hypothalamic (LH) MCH, Vglut2, and Hcrt neurons in *PS19* mice during simultaneous EEG/EMG monitoring (Fig. 7).

**Figure 7.**
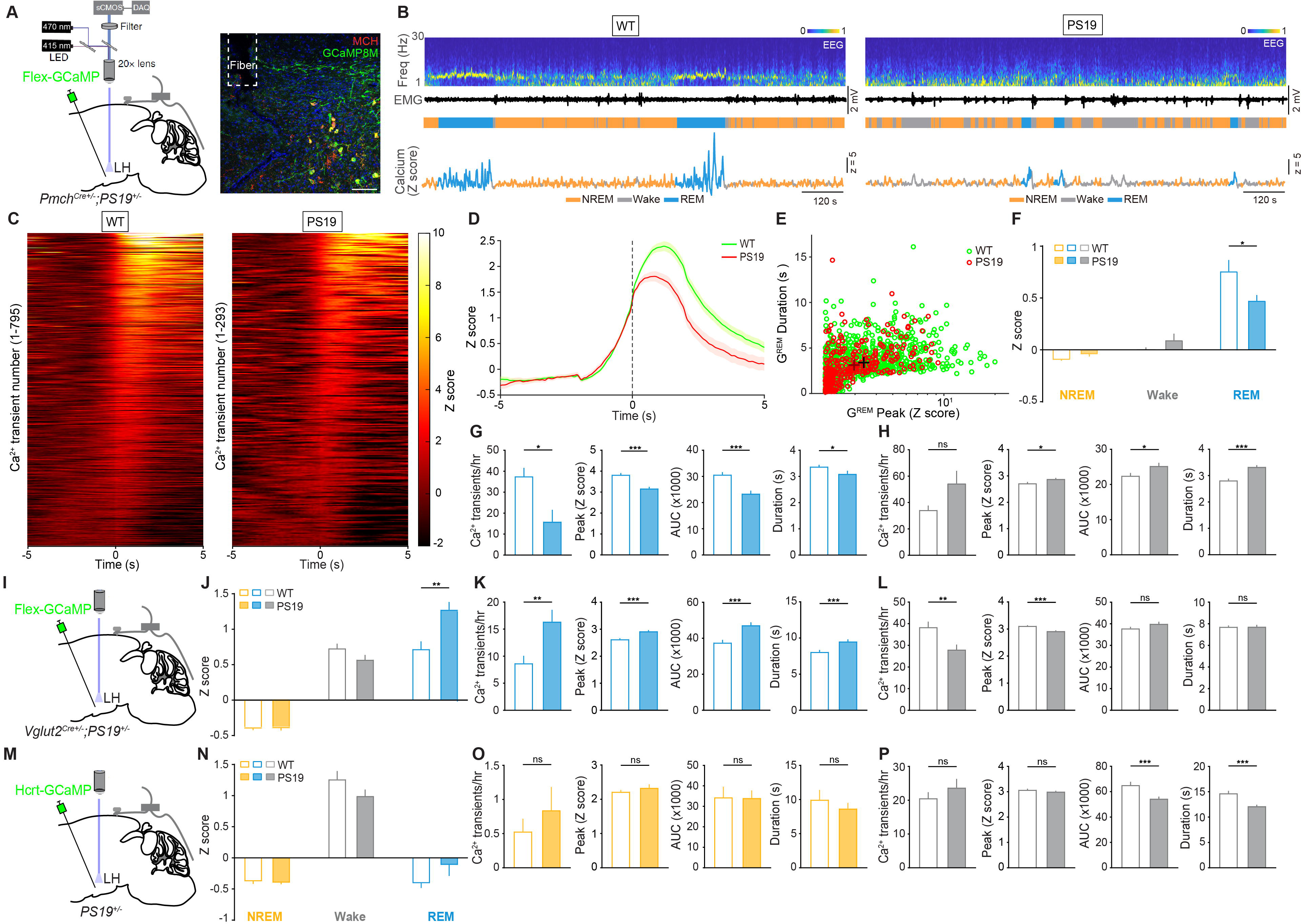
*PS19* mice show impaired state-dependent calcium activity in lateral hypothalamic sleep/wake neurons. (A) Left, schematic showing fiber photometry recording from lateral hypothalamic MCH neurons. AAV-Flex-jGCaMP8M was injected into the LH of *Pmch^Cre+/-^*;*PS19^+/-^* mice, and calcium signals were recorded through an implanted optical fiber using 470-nm excitation and 415-nm isosbestic control. Right, representative fluorescence image showing GCaMP8M expression in MCH neurons in the LH. Green, GCaMP8M; red, MCH. Dashed box indicates the fiber track. (B) Representative recordings from WT and *PS19* mice. Shown from top to bottom are EEG power spectrogram, EMG trace, color-coded brain states, and z-scored calcium signals recorded from LH MCH neurons. Yellow, NREM; gray, wake; blue, REM. Scale bars, as indicated. (C) Heatmaps of REM-associated MCH calcium transients in WT and *PS19* mice. Each row represents one calcium transient aligned to the transient peak. Color scale indicates z-scored calcium activity. (D) Average REM-associated MCH calcium transient traces from WT and *PS19* mice. Traces are aligned to the transient peak. Green, WT; red, *PS19*. Shading indicates ±SEM. (E) Relationship between REM-associated MCH calcium transient peak amplitude and duration in WT and *PS19* mice. Each dot represents one calcium transient. Green, WT; red, *PS19*. (F) Average MCH calcium activity (Z Score) during NREM sleep, wake, and REM sleep in WT and *PS19* mice. Yellow, NREM; gray, wake; blue, REM. n = 6-7 mice per group; Bars, mean ± SEM. *PS19* mice showed reduced REM-associated MCH calcium activity compared with WT mice. (G) Quantification of REM-associated MCH calcium transients in WT and *PS19* mice, including calcium transient frequency, peak amplitude, AUC, and duration. Bars, mean ± SEM. (H) Quantification of wake-associated MCH calcium transients in WT and *PS19* mice, including calcium transient frequency, peak amplitude, AUC, and duration. Bars, mean ± SEM. (I) Schematic showing fiber photometry recording from LH Vglut2 neurons. AAV-Flex-jGCaMP8M was injected into the LH of *Vglut2^Cre+/-^*;*PS19^+/-^* mice, and calcium activity was recorded during spontaneous sleep-wake behavior. (J) Average LH Vglut2 calcium activity (Z Score) during NREM sleep, wake, and REM sleep in WT and *PS19* mice. Orange, NREM; gray, wake; blue, REM. n = 7-11 mice per group; Bars, mean ± SEM. (K) Quantification of REM-associated LH Vglut2 calcium transients in WT and *PS19* mice, including calcium transient frequency, peak amplitude, AUC, and duration. Bars, mean ± SEM. (L) Quantification of wake-associated LH Vglut2 calcium transients in WT and *PS19* mice, including calcium transient frequency, peak amplitude, AUC, and duration. Bars, mean ± SEM. (M) Schematic showing fiber photometry recording from LH Hcrt neurons using an AAV-Hcrt-GCaMP6s strategy in *PS19* mice. (N) Average LH Hcrt calcium activity (Z Score) during NREM sleep, wake, and REM sleep in WT and *PS19* mice. Yellow, NREM; gray, wake; blue, REM. n = 8-9 mice per group; Bars, mean ± SEM. (O) Quantification of NREM-associated LH Hcrt calcium transients in WT and PS19 mice, including calcium transient frequency, peak amplitude, AUC, and duration. Bars, mean ± SEM. (P) Quantification of wake-associated LH Hcrt calcium transients in WT and PS19 mice, including calcium transient frequency, peak amplitude, AUC, and duration. Bars, mean ± SEM. Asterisks indicate significant differences between WT and *PS19* mice. Statistical significance for all comparisons was determined by Welch’s t-test. *P < 0.05; **P < 0.01; ***P < 0.001; ns, not significant.

To measure MCH neuronal activity, a Cre-dependent GCaMP8m virus (AAV-hSyn-FLEX-jGCaMP8m) was injected into the LH of *Pmch^Cre+/-^*;*PS19^+/-^*mice and their WT littermates (*Pmch^Cre+/-^*;*PS19^-/-^*) (Fig. 7A), and calcium activity was recorded at 10-11 months of age. Representative recordings revealed robust brain-state-dependent MCH neuronal activity in both genotypes, with the highest calcium activity observed during REM sleep (G^REM^), intermediate activity during wakefulness (G^wake^), and the lowest activity during NREM sleep (G^NREM^, Fig. 7B). We found that the amplitude changes of GCaMP signals were reduced in *Pmch^Cre+/-^*;*PS19^+/-^*mice (Figs. 7C and D), suggesting that the threshold of MCH neuronal activity associated with sleep-to-wake transitions is lowered by tau pathology. Compared with WT littermates, G^REM^ was significantly reduced in *Pmch^Cre+/-^*;*PS19^+/-^*mice (Figs. 7E and F), consistent with the notion that pathological tau accumulation impairs the REM sleep-promoting and sleep-maintaining functions of MCH neurons in 11-month-old transgenic mice. In addition, the calcium transient number, peak amplitude, area under the curve (AUC), and duration of G^REM^ were all reduced in *Pmch^Cre+/-^*;*PS19^+/-^* mice (Fig. 7G), whereas the peak amplitude, averaged AUC, and duration of G^wake^ were increased (Fig. 7H). In contrast, LH Vglut2 neurons exhibited an opposite pattern of activity changes. *Vglut2^Cre+/-^; PS19^+/-^* mice showed increased average activity, calcium transient number, peak amplitude, AUC, and duration of G^REM^ (Figs. 7J and K), along with decreased calcium transient number and peak amplitude of G^wake^ (Fig. 7L). For Hcrt neurons (Figs. 7M and N), no genotype differences were observed in G^NREM^ (Fig. 7O). However, the AUC and duration of G^wake^ were decreased in *PS19* mice (Fig. 7P). Together, these findings suggest that extensive pathological tau accumulation markedly impairs the REM sleep-promoting and sleep-maintaining functions of MCH neurons, profoundly alters the activity of Vglut2 neurons, and exerts comparatively milder effects on Hcrt neurons in the LH.

### Mutant P301L tau expression in MCH neurons induces sleep disturbances

Pathological tau is ubiquitously expressed throughout neuronal populations in the LH of *PS19* mice (Fig. 6), suggesting that the reduced MCH neuronal activity observed in these mice may not solely result from cell-autonomous effects. To directly determine whether tau pathology within MCH neurons is sufficient to impair neuronal activity and disrupt sleep, we used an AAV-mediated approach to selectively express mutant tau (P301L) in MCH neurons of WT (*Pmch^Cre^*) mice (Fig. 8A). In *Pmch^Cre^* mice injected with AAV-CMV-DIO-tau(P301L), we observed a significant loss of MCH neurons accompanied by robust pathological phosphorylated tau (pTau) expression compared with control mice injected with AAV-CMV-DIO-mCherry (Fig. 8B). Importantly, mutant tau-expressing mice also exhibited reduced NREM and REM sleep, along with increased wakefulness during the light phase, relative to mCherry-injected control mice (Fig. 8C). Using fiber photometry recordings, we next tested whether mutant tau affects MCH neuron activity. *Pmch^Cre^*mice at 3∼5 months of age were unilaterally injected into the LH with ∼700 nL of mixed AAV-CMV-DIO-tau (P301L)-mCherry or AAV-CMV-DIO-mCherry and AAV-hSyn-FLEX-jGCaMP8m. Fiber photometry recordings showed that mutant tau did not affect the overall z score of calcium signals (Fig. 8F), but significantly decreased the peak, AUC and duration of G^REM^ (Fig. 8G), and increased the peak of G^wake^ (Fig. 8H).

**Figure 8.**
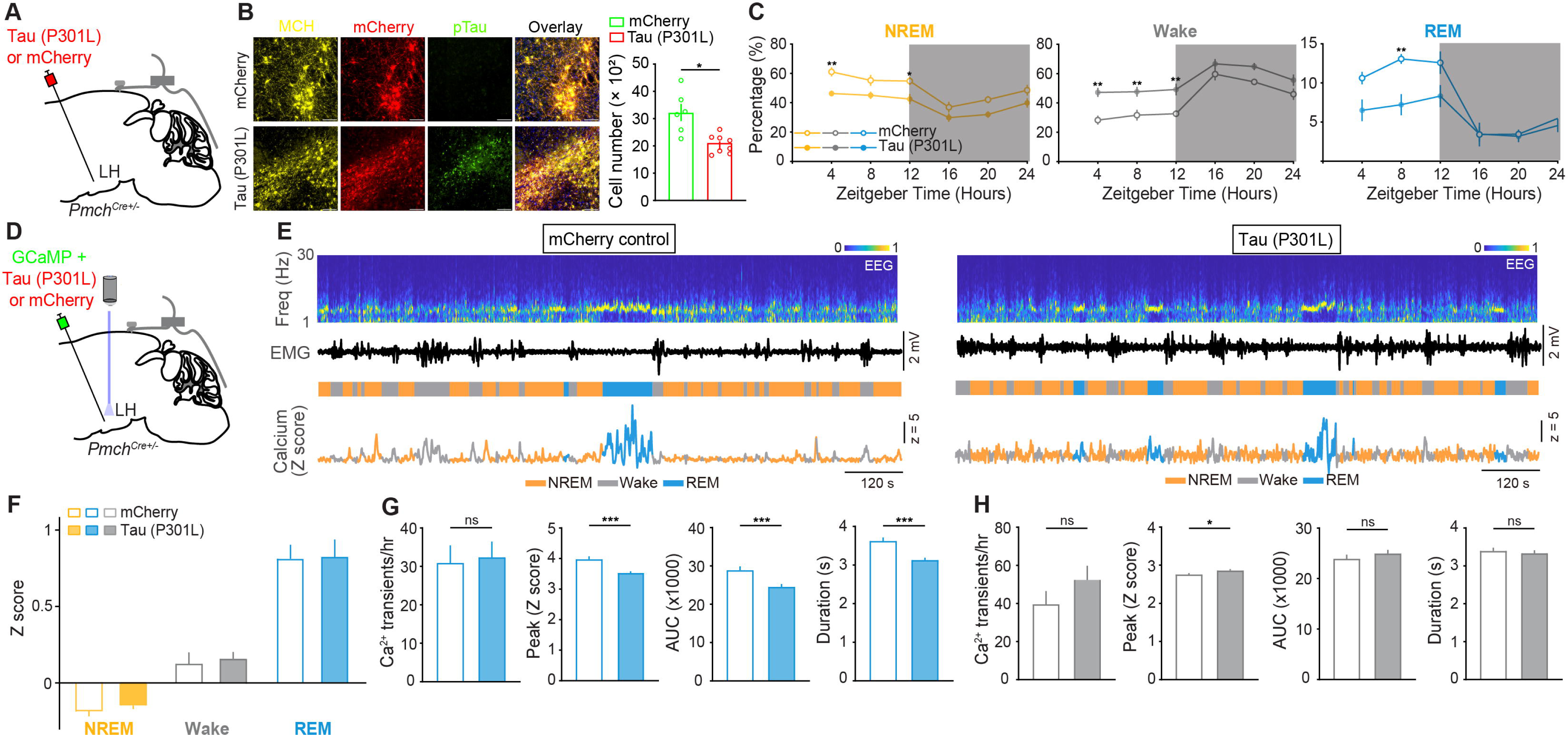
MCH neuron-specific Tau(P301L) expression disrupts sleep and impairs REM-associated MCH calcium activity. (A) Schematic showing viral expression of Tau(P301L) or mCherry control in MCH neurons. A Cre-dependent viral vector expressing Tau(P301L)-mCherry or mCherry was injected into the LH of *Pmch^Cre^* mice. (B) Left, representative immunofluorescence images showing MCH neurons, mCherry expression, and phosphorylated tau staining in the LH of mCherry control and Tau(P301L)-expressing mice. Yellow, MCH; red, mCherry; green, pTau. Overlay images show the spatial relationship between viral expression, MCH-positive neurons, and pTau signal. Scale bar, as indicated. Right, quantification of MCH-positive cell number in mCherry control and Tau(P301L)-expressing mice. Each circle represents one sample. N = 6-8 mice per group; Bars, mean ± SEM. Tau(P301L) expression significantly reduced the number of MCH-positive neurons compared with mCherry control. (C) Percentage of time spent in NREM sleep, wake, and REM sleep across the 24-h light/dark cycle in mCherry control and Tau(P301L)-expressing mice. Data are plotted as a function of Zeitgeber time. Gray shading indicates the dark phase. Orange, NREM; gray, wake; blue, REM. Open circles, mCherry control; filled circles, Tau(P301L). Error bars, ±SEM. Tau(P301L)-expressing mice showed reduced NREM sleep, increased wakefulness, and reduced REM sleep at selected Zeitgeber time points. (D) Schematic showing fiber photometry recording from LH MCH neurons following expression of GCaMP together with Tau(P301L) or mCherry control. Calcium signals were recorded through an implanted optical fiber during spontaneous sleep-wake behavior. (E) Representative recordings from mCherry control and Tau(P301L)-expressing mice. Shown from top to bottom are EEG power spectrogram, EMG trace, color-coded brain states, and z-scored calcium signals recorded from LH MCH neurons. Yellow, NREM; gray, wake; blue, REM. Scale bars, as indicated. (F) Average MCH calcium activity (Z Score) during NREM sleep, wake, and REM sleep in mCherry control and Tau(P301L)-expressing mice. Yellow, NREM; gray, wake; blue, REM. n = 6-11 mice per group; Bars, mean ± SEM. (G) Quantification of REM-associated MCH calcium transients in mCherry control and Tau(P301L)-expressing mice, including calcium transient frequency, peak amplitude, AUC, and duration. Bars, mean ± SEM. Tau(P301L)-expressing mice showed reduced REM-associated calcium transient peak, AUC, and duration compared with mCherry control. (H) Quantification of wake-associated MCH calcium transients in mCherry control and Tau(P301L)-expressing mice, including calcium transient frequency, peak amplitude, AUC, and duration. Bars, mean ± SEM. Tau(P301L)-expressing mice showed altered wake-associated calcium transient peak amplitude, whereas transient frequency, AUC, and duration were not significantly changed. Asterisks indicate significant differences between mCherry control and Tau(P301L)-expressing mice. Statistical significance for sleep-wake state percentage in (C) was determined by two-way ANOVA followed by post hoc multiple-comparisons tests. Statistical significance for MCH-positive cell number in (B), average calcium activity in (F), and calcium transient analyses in (G and H) was determined by Welch’s t-test. *P < 0.05; **P < 0.01; ***P < 0.001; ns, not significant.

### MCH neurons retain the capacity to regulate sleep at late stages of tauopathy

The above photometry recordings suggest that MCH neurons undergo persistent functional impairment in response to tau pathology (Figs. 7 and 8). We next used optogenetic and chemogenetic approaches to determine whether these neurons still retain the capacity to positively regulate sleep at late stages of tauopathy (Fig. 9). Cre-inducible AAV expressing ChR2-eYFP (AAV-DIO-hChR2(H134R)-eYFP) was unilaterally injected into the LH of *Pmch^Cre+/-^*;*PS19^+/-^*mice and their WT (*Pmch^Cr-^*;*PS19^-/-^*) littermates (Fig. 9A). Laser stimulation (2-4 mW, 20 Hz) was applied for 2 min/trial every 10-13 min during the light phase (Fig. 9B). Optogenetic activation of MCH neurons induced a robust increase in REM sleep and a concomitant decrease in NREM sleep in WT mice at 10–11 months of age (Fig. 9C). Notably, the laser-evoked changes in vigilance states were not significantly different between the WT and age-matched *Pmch^Cre+/-^*;*PS19^+/-^*mice (Fig. 9D), indicating that despite their functional impairments, MCH neurons retain the capacity to promote REM sleep at late stages of tauopathy.

**Figure 9.**
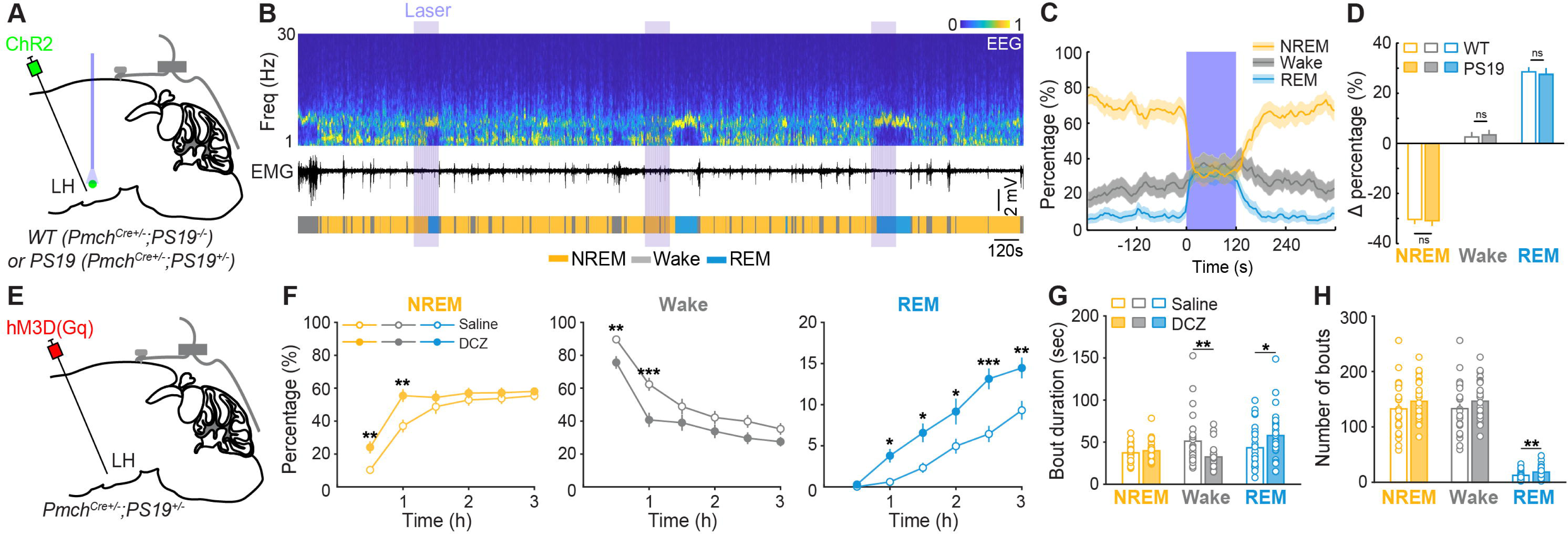
Activation of MCH neurons modulates sleep-wake states in aged *PS19* mice. (A) Schematic showing optogenetic activation of lateral hypothalamic MCH neurons. A Cre-dependent ChR2 virus was injected into the LH of WT (*Pmch^Cre+/-^*;*PS19^-/-^*) or *PS19* (*Pmch^Cre+/-^*;*PS19^+/-^*) mice, and an optical fiber was implanted above the LH for laser stimulation during sleep-wake recording. (B) Representative recording showing the effect of optogenetic stimulation of MCH neurons. Shown from top to bottom are EEG power spectrogram, EMG trace, and color-coded brain states. Purple shading indicates laser stimulation periods. Yellow, NREM; gray, wake; blue, REM. Scale bars, as indicated. (C) Percentage of time spent in NREM sleep, wake, and REM sleep before, during, and after laser stimulation. Blue shading indicates the laser stimulation period. Data are aligned to laser onset. Yellow, NREM; gray, wake; blue, REM. Lines and shading indicate mean ± SEM. (D) Quantification of laser-induced changes in the percentage of time spent in NREM sleep, wake, and REM sleep in WT and *PS19* mice. Δ percentage was calculated by comparing sleep-wake state occupancy before and during laser stimulation. Open bars, WT; filled bars, PS19. n = 6 mice per group; Bars, mean ± SEM. No significant genotype difference was detected in laser-induced changes in NREM sleep, wake, or REM sleep. (E) Schematic showing chemogenetic activation of MCH neurons. A Cre-dependent hM3D(Gq) virus was injected into the LH of *Pmch^Cre+/-^*;*PS19^+/-^* mice, and sleep-wake behavior was recorded following saline or DCZ administration. (F) Percentage of time spent in NREM sleep, wake, and REM sleep following saline or DCZ administration. Data are plotted over the 3-h recording period after injection. Orange, NREM; gray, wake; blue, REM. Open circles, saline; filled circles, DCZ. n = 6 mice per group; Error bars, ±SEM. Chemogenetic activation of LH MCH neurons increased NREM sleep and REM sleep while reducing wakefulness at selected time points. (G) Mean bout duration of NREM sleep, wake, and REM sleep following saline or DCZ administration. Each circle represents one sample. Open bars, saline; filled bars, DCZ. Bars, mean ± SEM. DCZ treatment increased wake and REM bout duration compared with saline. (H) Number of NREM, wake, and REM bouts following saline or DCZ administration. Each circle represents one sample. Open bars, saline; filled bars, DCZ. Bars, mean ± SEM. DCZ treatment reduced the number of REM bouts compared with saline. Asterisks indicate significant differences between groups or treatments. Statistical significance for sleep-wake state percentage following saline or DCZ administration in (F) was determined by two-way ANOVA followed by post hoc multiple-comparisons tests. Statistical significance for laser-induced changes in sleep-wake state percentage in (D), mean bout duration in (G), and bout number in (H) was determined by Welch’s t-test. *P < 0.05; **P < 0.01; ***P < 0.001; ns, not significant.

We then tested the effect of chemogenetic activation of MCH neurons on sleep-wake regulation in *PS19* mice at late stages of tauopathy injecting AAV-DIO-hM3D(Gq)-mCherry into the LH of *Pmch^Cre+/-^*;*PS19^+/-^*mice (Fig. 9E). Compared to saline injection, DREADD receptor agonist, deschloroclozapine dihydrochloride (DCZ)(*41*) injection, strongly increased NREM and REM sleep and decreased wakefulness in *Pmch^Cre+/-^*;*PS19^+/-^* mice at 10∼11 months of age (Fig. 9F). DCZ injection also improved sleep quality by increasing REM sleep bout duration, and decreasing wake bout duration (Fig. 9G), as well as increasing the number of REM sleep bouts (Fig. 9H).

We next tested whether enhancing MCH neuronal activity could ameliorate sleep deficits in *PS19* mice at late stages of tauopathy. To selectively activate MCH neurons, we injected a Cre-dependent excitatory DREADD virus (AAV-DIO-hM3D(Gq)-mCherry) into the LH of *Pmch^Cre+/-^*;*PS19^+/-^* mice (Fig. 9E). Administration (i.p.) of the DREADD agonist deschloroclozapine (DCZ), compared with saline treatment, robustly increased both NREM and REM sleep while reducing wakefulness in 10∼11-month-old *Pmch^Cre+/-^*;*PS19^+/-^* mice (Fig. 9F). In addition to increasing total sleep time, DCZ improved sleep architecture by prolonging REM sleep bout duration and shortening wake bout duration (Fig. 9G). DCZ treatment also increased the number of REM sleep bouts (Fig. 9H), indicating enhanced REM sleep expression. These findings demonstrate that acute activation of MCH neurons effectively restores sleep quantity and quality in *PS19* mice despite the presence of advanced tau pathology.

To evaluate the therapeutic potential of enhancing MCH neuronal activity for correcting disease-associated sleep disturbances, we examined the effects of repeated chemogenetic activation of MCH neurons on 24-hour sleep–wake architecture in 10∼11-month-old *Pmch^Cre+/-^*;*PS19^+/-^*mice (Fig. 10A). Mice expressing hM3D(Gq) in MCH neurons received bi-daily injection of either saline or DCZ at ZT0 and ZT12 for 3 consecutive days (Fig. 10B). Relative to saline treatment, repeated DCZ administration significantly increased both NREM and REM sleep while reducing wakefulness across the 24-hour recording period (Fig. 10C). These findings demonstrate that sustained activation of MCH neurons effectively restores sleep in *PS19* mice and suggest that dysfunction of the MCH system contributes directly to the sleep disturbances associated with tauopathy.

**Figure 10.**
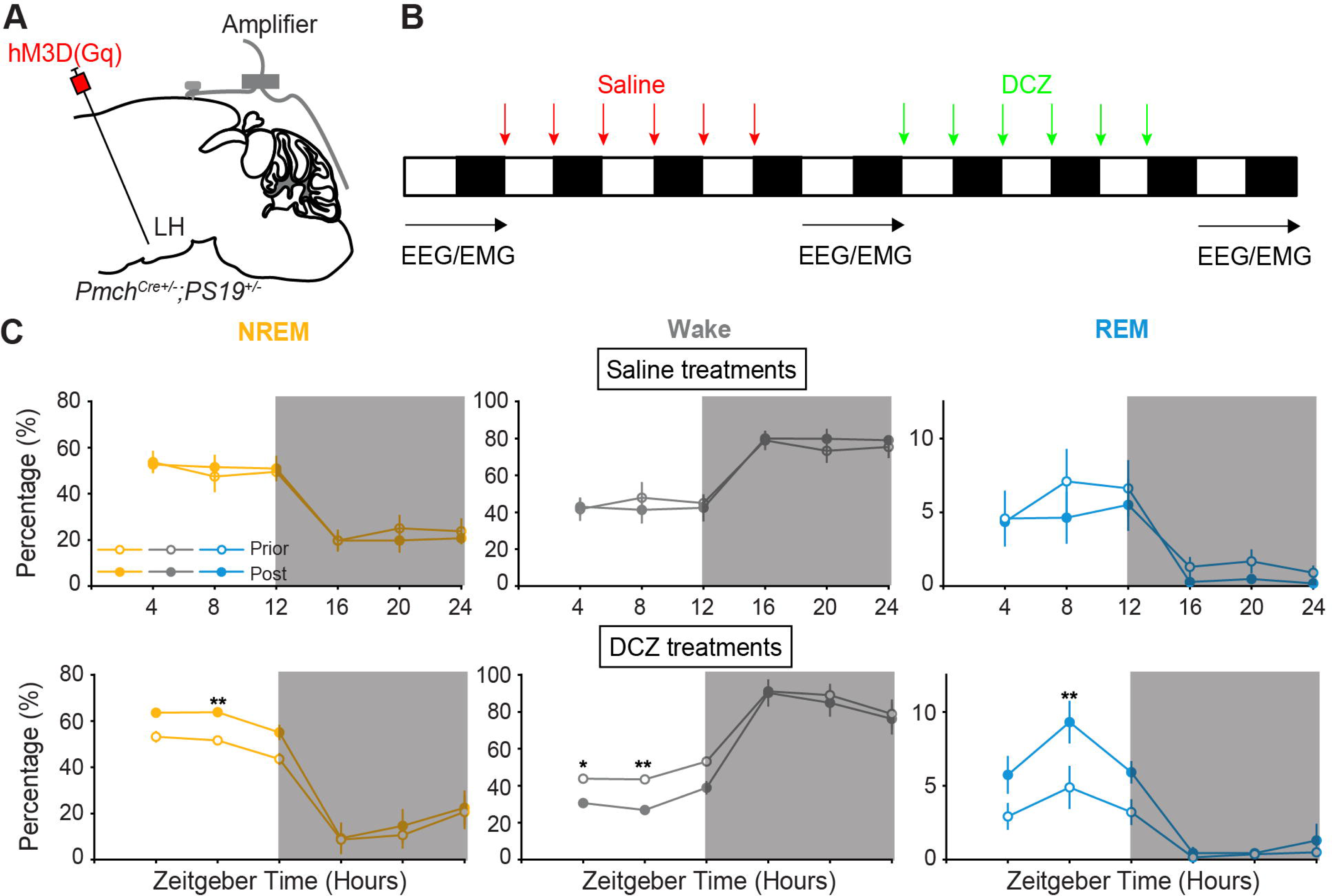
Repeated chemogenetic activation of MCH neurons restores sleep in aged *PS19* mice. (A) Schematic showing chemogenetic activation of MCH neurons during EEG/EMG recording. A Cre-dependent hM3D(Gq) virus was injected into the LH of *Pmch^Cre+/-^* ;*PS19^+/-^* mice, and EEG/EMG electrodes were implanted to monitor sleep-wake states. (B) Experimental timeline for repeated saline and DCZ treatments. EEG/EMG recordings were performed before and after repeated saline treatment and before and after repeated DCZ treatment. Red arrows indicate saline injections, and green arrows indicate DCZ injections. Black and white bars indicate the light/dark cycle. (C) Percentage of time spent in NREM sleep, wake, and REM sleep before and after repeated saline or DCZ treatments. Top, sleep-wake state distribution before and after saline treatment. Bottom, sleep-wake state distribution before and after DCZ treatment. Data are plotted as a function of Zeitgeber time. Gray shading indicates the dark phase. Yellow, NREM; gray, wake; blue, REM. Open circles, prior to treatment; filled circles, post treatment. n = 5 mice per group; Error bars, ±SEM. Repeated saline treatment produced minimal changes in sleep-wake state distribution, whereas repeated DCZ treatment decreased NREM sleep and REM sleep and increased wakefulness during the light phase at selected Zeitgeber time points. Asterisks indicate significant differences between prior and post treatment at individual Zeitgeber time points. Statistical significance for sleep-wake state percentage in (C) was determined by two-way ANOVA followed by post hoc multiple-comparisons tests. *P < 0.05; **P < 0.01.

## DISCUSSION

Sleep disturbances are among the earliest and most prevalent symptoms of AD and related tauopathies (*1, 2*), yet the neural mechanisms linking tau pathology to disrupted sleep remain poorly understood. In this study, we combined longitudinal EEG/EMG recordings, fiber photometry, histological analyses, and causal circuit manipulations to investigate how tau pathology affects sleep-regulatory circuits in the lateral hypothalamus. We demonstrate that tauopathy is associated with progressive impairments in sleep architecture and sleep homeostasis, degeneration of lateral hypothalamic MCH and Hcrt neurons, and selective dysfunction of MCH neurons during REM sleep. Importantly, despite substantial neuronal loss and reduced endogenous activity, surviving MCH neurons retain their sleep-promoting capacity, and their activation effectively restores sleep in tauopathy mice. Together, these findings identify dysfunction of the MCH system as a previously underappreciated mechanism contributing to sleep disturbances in tauopathy and suggest that MCH neurons represent a promising therapeutic target for treating sleep dysfunction in AD.

Previous clinical and experimental studies have established a strong association between sleep disturbances and tau pathology. Sleep disruption frequently precedes cognitive decline in AD patients (*5–7*), while sleep loss accelerates tau release, aggregation, and propagation (*42*). Conversely, tau pathology accumulates early in several brain regions involved in sleep-wake regulation (*3*). Consistent with these observations, we found that both *PS19* and *hTau* mice develop abnormalities in sleep regulation and EEG spectra, although the severity and temporal progression differed between models. *PS19* mice exhibited profound reductions in NREM and REM sleep, fragmented sleep architecture (Fig. 1), altered EEG oscillations (Fig. 2), and impaired homeostatic sleep responses at late disease stages (Fig. 5A). In contrast, *hTau* mice displayed milder and later-emerging phenotypes characterized by altered REM sleep regulation (Fig. 5B), impaired wake-state consolidation (Fig. 3), and reduced NREM slow-wave activity (Fig. 4). The presence of sleep abnormalities in two independent tauopathy models supports the conclusion that sleep dysfunction is a conserved consequence of pathological tau accumulation rather than a model-specific phenomenon.

One of the most notable findings of this study is the selective vulnerability of the MCH system to tau pathology. Histological analyses revealed degeneration of both MCH and Hcrt neurons in the lateral hypothalamus of *PS19* mice (Fig. 6). However, calcium photometry demonstrated that MCH neurons underwent substantially greater functional impairment than Hcrt neurons (Fig. 7). MCH neuronal activity during REM sleep was markedly reduced, accompanied by decreases in calcium transient frequency, amplitude, duration, and integrated activity. In contrast, Hcrt neurons exhibited relatively modest changes despite significant tau accumulation within the lateral hypothalamus. These findings are particularly intriguing given that MCH neurons are strongly implicated in REM sleep regulation and sleep homeostasis. The observed reduction in REM-associated MCH activity therefore provides a mechanistic explanation for the REM sleep deficits and impaired sleep recovery observed in tauopathy mice.

Our findings also provide evidence that tau pathology directly disrupts MCH neuronal function. In *PS19* mice, pathological tau is broadly expressed throughout the brain, raising the possibility that impaired MCH activity could arise indirectly through network dysfunction. To address this question, we selectively expressed mutant P301L tau in MCH neurons of otherwise healthy mice (Fig. 8). Cell-specific tau expression was sufficient to induce MCH neuronal loss, reduce REM-associated MCH activity, and produce sleep disturbances resembling those observed in *PS19* mice. These findings indicate that tau pathology exerts cell-autonomous effects on MCH neurons and suggest that intrinsic dysfunction of the MCH system contributes directly to sleep abnormalities during disease progression. Given that MCH neurons also regulate hippocampal activity and memory processes (*33, 43*), it will be important in future studies to determine whether MCH dysfunction contributes not only to sleep disturbances but also to cognitive deficits associated with tauopathy.

An unexpected finding was that surviving MCH neurons retained substantial functional capacity despite profound structural and physiological impairments. Both optogenetic and chemogenetic activation robustly promoted sleep in *PS19* mice at advanced stages of disease (Fig. 9). Acute chemogenetic activation increased NREM and REM sleep, improved sleep architecture, and reduced wakefulness, whereas repeated activation restored sleep across the 24-hour cycle (Fig. 10). These observations indicate that the sleep-promoting circuitry downstream of MCH neurons remains largely intact despite extensive tau pathology. More broadly, they suggest that disease-associated sleep disturbances may arise in part from reduced activity within surviving MCH neurons rather than irreversible loss of circuit function. This distinction has important therapeutic implications because it implies that enhancing the activity of remaining MCH neurons could compensate for neuronal loss and restore sleep even after pathology has become established.

The mechanisms responsible for MCH neuronal dysfunction remain to be determined. Tau pathology may impair intrinsic excitability, disrupt synaptic connectivity, alter neurotransmitter release, or interfere with cellular homeostatic processes required for state-dependent activation. The reciprocal changes observed in MCH and Vglut2 neuronal activity (Fig. 8) raise the possibility that tau pathology also alters local hypothalamic circuit interactions that regulate sleep-wake transitions. Furthermore, because MCH neurons integrate homeostatic sleep pressure and coordinate REM sleep expression (*23–26*), reduced MCH activity may contribute simultaneously to impaired sleep maintenance, reduced REM sleep, and deficits in sleep rebound following deprivation. Future studies combining circuit mapping, slice electrophysiology, and transcriptomic profiling will be valuable for defining the molecular and network mechanisms underlying MCH vulnerability during tauopathy.

Several limitations should be considered. Although our data establish a causal role for MCH dysfunction in sleep disturbances, additional sleep-regulatory populations are also affected by tau pathology and likely contribute to disease phenotypes. The degeneration of Hcrt neurons and altered activity of LH glutamatergic neurons observed here suggest that sleep dysfunction results from broader network-level abnormalities. In addition, while restoration of sleep by MCH activation strongly supports a therapeutic role for this circuit, the long-term consequences of enhancing MCH signaling on cognitive decline, tau pathology, and neurodegeneration remain unknown. Determining whether sustained correction of sleep disturbances can slow or halt disease progression represents an important direction for future investigation.

In summary, our study identifies the lateral hypothalamic MCH system as a critical target of tau pathology and reveals a mechanistic link between MCH dysfunction and sleep disturbances in tauopathy. We demonstrate that tau pathology causes degeneration and functional impairment of MCH neurons, that selective tau expression within MCH neurons is sufficient to induce sleep abnormalities, and that enhancing MCH activity restores sleep despite advanced disease pathology. These findings provide new insight into the circuit mechanisms underlying sleep dysfunction in AD and related tauopathies and highlight the MCH system as a potential therapeutic target for improving sleep and potentially modifying disease progression.

## MATERIALS AND METHODS

### Animals

*Vglut2^Cre^*, *PS19* and *hTau* (Jackson stock no: 016963, 008169, and 005491 respectively) were obtained from Jackson Laboratory. *Pmch^Cre^* mice were obtained from SRI International. Mice were housed in 12 hrs light-dark cycle (lights on 06:00 am and off at 06:00 pm) with free access to food and water. Experimental mice were allowed ∼3 weeks for postoperative recovery and acclimation to the recording apparatus before data collection. All procedures were approved by the Animal Care and Use Committees of the University of Nebraska Medical Center.

### Surgeries

Mice were anesthetized with 1.5% isoflurane and placed on a stereotaxic frame (RWD Life Science). Body temperature was kept stable throughout the procedure using a heating pad. After asepsis, the skin was incised to expose the skull, and the overlying connective tissue was removed. To implant EEG and EMG recording electrodes, two stainless steel screws were inserted into the skull 1.5 mm from midline and 1.5 mm anterior to the bregma, and two others were inserted 3 mm from midline and 3.5 mm posterior to the bregma. Two EMG electrodes were inserted into the neck musculature. Insulated leads from the EEG and EMG electrodes were soldered to a pin header, which was secured to the skull using dental cement.

For fiber photometry and optogenetic experiments, a craniotomy (0.5-1mm diameter) was made on top of the LH, and 0.4-0.5 µl virus was unilaterally injected into the LH (AP -1.4 mm, ML 1.2 mm, DV -4.6 mm) using Nanoject II (Drummond Scientific) via a micro pipette. Optic fibers (0.2 mm diameter; Thorlabs, MBF Bioscience or RWD Life Science) were implanted into the target region with the tip 0.2 mm above the virus injection site two weeks after viral injection. Dental cement was applied to cover the exposed skull completely and to secure the implants for EEG and EMG recordings to the screws. Data from animals used in experiments were excluded based on histological criteria that included injection sites, virus expression and optical fiber placement. Only animals with injection sites and optic fiber placement in the region of interest were included. For chemogenetic experiments, ∼0.3 µl AAV2-EF1α-DIO-hM3D(Gq)-mCherry (bilateral) was injected into the target region.

### EEG/EMG Recordings

Behavioral experiments were carried out in home cages placed in sound-attenuating boxes between 6:00 am and 6:00 pm (light phase) and between 8:00 pm and 12:00 am (dark phase). EEG and EMG electrodes were connected to flexible recording cables via a mini-connector. Recordings started after 20-30 min of habituation. The signals were recorded with an Open Ephys Acquisition Board (bandpass filter, 0.1-1,000 Hz; sampling rate, 1,000 Hz).

For sleep scoring, EEG and EMG spectral analysis was carried out using fast Fourier transform (FFT). We first calculated the power spectrum for the EEG and EMG signals using 5 s sliding windows, sequentially shifted by 2.5 s increments. Activity states were visually classified as wake, rapid eye movement sleep (REM) or non-REM (NREM) in 2.5 sec epochs. Spectral analysis was carried out using fast Fourier transform, and brain states were classified in 2.5 sec epoch using Accusleep(*44*). Briefly, wakefulness was characterized by desynchronized EEG and high EMG activity. NREM sleep was characterized by synchronized EEG with high-amplitude, low-frequency (1 to 4 Hz) activity and low EMG activity. REM sleep was characterized by high EEG power at theta frequencies (6 to 9 Hz) and low EMG activity. To train the neural network in Accusleep, a subset of data was manually scored and used to guide automated classification. A bout was defined as having two consecutive epochs (i.e. 4 sec) in the same state. Periods of seizure activity as well as any EEG/EMG activity artifacts associated with motor behavior were removed from analysis. Each recording was then visually inspected and manually corrected to resolve discrepancies.

### Fiber Photometry Recordings and Analysis

Calcium dynamics and EEG/EMG were recorded for ∼2-3 hours across sleep/wake states. TTL pulses were recorded simultaneously with EEG/EMG during photometry recordings. Fluorescent excitation was delivered using interleaved triggering of 470-nm and 415-nm LEDs (10-Hz trigger rate for each channel), and emission was measured with a Complementary Metal Oxide Semiconductor camera (FP3002, MBF Bioscience).

The 415-nm channel was used to correct for nonspecific, calcium-independent fluorescence fluctuations. For each recording session, 470- and 415-nm signals were first smoothed using a moving-average filter to correct for baseline drift caused by photobleaching. Slow baseline drift was estimated using adaptive iteratively reweighted penalized least squares and subtracted from each channel. The baseline-corrected 415-nm signal was then linearly fitted to the baseline-corrected 470-nm signal using least-squares regression, and the fitted 415-nm component was subtracted from the 470-nm signal to obtain the corrected fluorescence signal (*45*). The corrected signal was converted to z scores using the mean and standard deviation of the analyzed recording period.

Calcium transients were detected from the z-scored signal using a threshold-based method. Candidate transients were identified when the z-scored signal exceeded the mean + 2 standard deviation. For each transient, the peak amplitude was defined as the maximum z score between threshold-crossing onset and offset. Transient boundaries were further estimated relative to the local baseline, and transient AUC was calculated as the summed absolute difference between the z-scored signal and its estimated baseline within the transient boundaries. Transients were classified as REM, wake, NREM, or microarousal according to the behavioral state at the transient peak.

### Optogenetic Manipulation

The mice were habituated for two days before any behavioral testing, and in each testing session they were habituated for > 30 min before data collection. Each optic fiber was attached through an FC/PC adaptor to a 473-nm blue laser diode (Laserglow Technologies). For optogenetic activation, each trial consisted of a 20 Hz pulse train (10 ms per pulse, 2-4 mW at fiber tip) lasting for 120 s. In each optogenetic manipulation experiment, inter-trial interval was chosen randomly from a uniform distribution between 10 and 13 min. Each experimental session lasted for 3 hrs, and each animal was tested in 6-8 sessions.

### Chemogenetic Manipulation

Saline (0.9% NaCl) or DCZ dissolved in saline was injected intraperitoneally (i.p.) into the corresponding Cre mice expressing hM3D(Gq) in the target region. DCZ dose was 0.3 mg/kg body weight. For acute activation, each recording session started immediately after injection. In each test set, mice were administered with saline on the first day and DCZ on the second day. Each mouse was subjected to 3-4 test sets, and the data were averaged across all sets. For repeated activation, mice received bi-daily injections at ZT0 and ZT12 for 3 consecutive days, first with saline and subsequently with DCZ (a 3-day washout period). Continuous EEG/EMG recordings were collected during baseline conditions and following each treatment period.

### Histology

Mice were deeply anaesthetized and transcardially perfused with 0.1M PBS followed by 4% paraformaldehyde (w/v) in PBS. For fixation, brains were kept overnight in 4% paraformaldehyde. For cryoprotection, brains were placed in 30% sucrose (w/v) in PBS solution for 36-48 hrs. After embedding and freezing, brains were sectioned into 30∼50 µm coronal slices using a cryostat (RWD Life Science). For immunofluorescence staining, frozen brain sections were baked at 55 °C for 1 h and then washed in 1× PBS for 10 min three times to remove residual OCT compound. Heat-induced epitope retrieval was performed by incubating the sections in antigen retrieval buffer at 80 °C for 30 min in a water bath, followed by gradual cooling to room temperature for approximately 2 h. Sections were washed in PBS for 5 min three times and then permeabilized and blocked in blocking buffer containing 5% normal goat serum and 0.3% Triton X-100 in PBS for 1 h at room temperature. Without additional washing, sections were incubated with primary antibodies diluted in blocking buffer at 4 °C for 36 h. Primary antibodies included anti-MCH antibody (1:500), anti-Hcrt antibody (1:250), anti-phospho-Tau (AT8) antibody (1:500), anti-mCherry antibody (1:500), and anti-GFP antibody (1:500). Sections were then washed in PBS for 10 min three times and incubated with species-appropriate fluorescent secondary antibodies diluted at 1:500 in blocking buffer for 2 h at room temperature protected from light. After incubation, sections were washed in PBS for 20 min three times, counterstained with Hoechst (1:10,000) for 5 min, washed again in PBS, and mounted with antifade mounting medium. Fluorescence images were taken using a Nikon A1R confocal microscope and an Evident Scientific VS-200 Slide Scanner.

### Statistics

Statistical analyses were performed using MATLAB, SigmaPlot, and GraphPad Prism. All statistical tests were two-sided. For long-term sleep recordings and chemogenetic experiments, one-way or two-way repeated-measures ANOVA was used to assess the effects of genotype, treatment, time, or their interactions, as appropriate. When ANOVA indicated a significant effect or interaction, post hoc multiple-comparisons tests were performed to compare relevant groups or time points. Welch’s t-test was used for independent two-group comparisons when appropriate. Statistical significance was defined as P < 0.05.

For optogenetic experiments, the 95% confidence intervals (CI) for brain state probabilities were calculated using a bootstrap procedure: For an experimental group of *n* mice, with mouse *i* comprising *m_i_* trials, we repeatedly resampled the data by randomly drawing for each mouse *m_i_* trials (random sampling with replacement; each trial is a 600 s segment, containing 240 s before, 120 s during, and 240 s after the laser stimulation period). For each of the 10,000 iterations, we recalculated the mean probabilities for each brain state across the *n* mice. The lower and upper confidence intervals were then extracted from the distribution of the resampled mean values. To test whether a given brain state is significantly modulated by laser stimulation, we calculated for each bootstrap iteration the difference between the mean probabilities during laser stimulation and the preceding period of identical duration. To compare the degree of laser-induced brain state changes between two groups of animals, in each iteration we resampled the data for each group separately. The laser effect (difference in brain state probability between the pre-laser and laser periods) was computed for each group, and the difference between the two groups was computed. The P value was determined from the distribution of the difference in 10,000 iterations.

## ACKNOWLEDGEMENTS

This work was supported by the National Institutes of Health (NIH) under Award Numbers P20 GM130447 (PZ) and T32 AG076407 (WDM), and CurePSP under Award Number 703-2024-07 (PZ). We thank Dr. Thomas Kilduff at SRI International for generously providing the *Pmch^Cre^*mice used in this study.

## AUTHOR CONTRIBUTIONS

K.P. and P.Z. designed the study and wrote the manuscript. K.P. performed most of the experiments. S.G., A.K., L.X., W.M., Q.L. and D. J. performed some EEG/EMG recordings and histological studies. R.Y. wrote part of programs for data analysis. P.Z. supervised all aspects of the work.

